# Activation of VTA GABA Neurons Disrupts Reward Seeking by Altering Temporal Processing

**DOI:** 10.1101/2020.08.18.256073

**Authors:** Andrea K. Shields, Mauricio Suarez, Ken T. Wakabayashi, Caroline E. Bass

**Affiliations:** Department of Pharmacology and Toxicology, Jacobs School of Medicine and Biomedical Sciences, University at Buffalo, SUNY, Buffalo, New York, 14214; Clinical Research Institute on Addictions, University at Buffalo, State University of New York, Buffalo, New York 14203; Department of Psychology, University of Nebraska-Lincoln, 1220 T. Street, Lincoln, NE 68588

**Keywords:** GABA, ventral tegmental area, chemogenetics, reinforcement, rat, fixed interval

## Abstract

The role of ventral tegmental area (VTA) dopamine in reward, cue processing, and interval timing is well characterized. Using a combinatorial viral approach to target activating DREADDs (<u>D</u>esigner <u>R</u>eceptors <u>E</u>xclusively <u>A</u>ctivated by <u>D</u>esigner <u>D</u>rugs, hM3D) to GABAergic neurons in the VTA of male rats, we previously showed that activation disrupts responding to reward-predictive cues. Here we explored how VTA GABA neurons influence the perception of time in two fixed interval (FI) tasks, one where the reward or interval is not paired with predictive cues (Non-Cued FI), and another where the start of the FI is signaled by a constant tone that continues until the rewarded response is emitted (Cued FI). Under vehicle conditions in both tasks, responding was characterized by “scalloping” over the 30s FI, in which responding increased towards the end of the FI. However, when VTA GABA neurons were activated in the Non-Cued FI, the time between the end of the 30s interval and when the rats made a reinforced response increased. Additionally, post-reinforcement pauses and overall session length increased. In the Cued FI task, VTA GABA activation produced erratic responding, with a decrease in earned rewards. Thus, while both tasks were disrupted by VTA GABA activation, responding that is constrained by a cue was more sensitive to this manipulation, possibly due to convergent effects on timing and cue processing. Together these results demonstrate that VTA GABA activity disrupts the perception of interval timing, particularly when the timing is set by cues.

## I. Introduction

The role of the ventral tegmental area (VTA) in motivation and drug seeking behaviors has been well established [1-9]. The VTA is comprised primarily of dopamine (DA) neurons (∼60-70%), with the remaining neurons being γ-aminobutyric acid (GABA, 30-35%) and glutamate (∼1-2%) [10-12]. DA has a clear role in motivation, drug seeking [13], and cue-processing [14-17]. DA also contributes to interval timing, which occurs in the range of seconds to minutes, [18-22] where it plays an integral role in regulating the temporal control over behavior [18-20, 23-25]. For example, DA agonists, and pharmacological manipulations that increase DA release, accelerate the internal clock causing subjects to underestimate the interval time and respond sooner in a timed interval task [18, 21, 26] whereas decreasing dopamine produces the opposite effect [24].

The ability to time events has an important role in successfully performing operant tasks. Temporal control of behavior can be evaluated through the use of fixed interval (FI) schedules of reinforcement [18, 19, 27]. Using this model, selective knockdown of VTA tyrosine hydroxylase decreases DA, leading to a disruption of interval timing and reduces responding at the end of the interval [19]. In other areas of the mesocorticolimbic circuit, drugs of abuse such as amphetamine, and other DA agonists [25, 27] that increase DA in the nucleus accumbens (NAc) accelerate the internal clock. Moreover, microinfusion of the D1 receptor antagonist SCH23390 in the prefrontal cortex (PFC) disrupts interval timing by flattening response curves from the typical scalloped pattern of responding exhibited in control subjects [19]. Experiments using fast-scan cyclic voltammetry [18] revealed that the DA concentration in the nucleus accumbens (NAc) is inversely proportional to the perceived duration of the interval, and pharmacological enhancement of NAc DA decreased the latency to respond within a fixed interval. Thus, VTA DA acts as a signal controlling the perceived length of the interval, and treatments that alter DA kinetics in the NAc influence the perception of time. Several studies suggest that VTA GABA strongly impacts DA signaling, and in turn, the motivating and rewarding properties of drugs of abuse. GABA regulation of VTA DA occurs through local inhibition, likely by GABA interneurons, and GABA projections to other brain regions including the NAc, PFC, and amygdala [28-30]. Functionally, local VTA GABA neurons increase firing in response to an aversive stimulus (e.g. foot shock), which inhibits VTA DA firing via GABA_A_ receptors [31]. VTA GABA activation has also been shown to induce conditioned place aversion [32, 33]. Yet others have shown that while VTA GABA activation decreases reward consummatory behaviors, it does not impact cue-induced reward seeking [32]. In contrast, we recently demonstrated that VTA GABA activation greatly attenuates reward seeking in an operant task heavily dependent on cue responding, while chemogenetic activation of the VTA GABA terminals in the NAc had no effect [34]. Other studies have shown that VTA GABA activation promotes sleep [35] and head orienting towards desired spatial targets [36]. Thus, it is becoming clear that VTA GABA neurons contribute to the regulation and control of a myriad of motivated behaviors.

In the current study, we evaluated the role of VTA GABA neurons in interval timing for a palatable reinforcer (e.g. sucrose) during FI schedules. We hypothesized that activating VTA GABA neurons would disrupt FI responding, resulting in an overestimation of the interval time and shifting the scalloped responding pattern to later in the interval. Further we predicted that the disruption would be greater in a timed operant task in which the start of the intervals was signaled by a cue. To that end, two separate FI procedures were used: in the first there was no cue to signal the start of the interval or the end when sucrose was available (Non-Cued FI), while in the second, a tone indicated the start of a 30s fixed interval, and continued until a rewarded response was made after the 30s FI had elapsed (Cued FI).

## 2. Materials and Methods

### 2.1. Subjects

Experimentally naϊve adult male Long Evans rats (N=32) were obtained from Envigo RMS Incorporated (Indianapolis, IN), individually housed in transparent polycarbonate cages (20 cm x 46 cm x 20 cm) and maintained at 80-85% of free-feeding weight while potable tap water was available *ad libitum*. Rats were maintained on a 12:12 light cycle (lights on at 3PM) in a temperature and humidity-controlled room. Experimental procedures were conducted between 12PM to 2PM; all procedures were in accordance to the ARRIVE guidelines, NIH Guide for Care and Use of Laboratory Animals and approved by the Institutional Animal Care and Use Committee of the University at Buffalo, SUNY.

### 2.2. Adeno-associated virus (AAV) infusion

Rats were given five days to acclimate to the facility before stereotaxic surgery. Anesthesia was induced with a mix of 33 mg/kg ketamine and 10 mg/kg xylazine intraperitoneally (i.p.) and maintained with 1-2% isoflurane, delivered through a nosecone affixed to the stereotaxic frame. An incision was made at the midline of the scalp, and holes were drilled through the skull above the VTA. A glass micropipette (Drummond Scientific Company, Broomall, PA), affixed to a Nanoject III (Drummond Scientific Company, Broomall, PA) pump was used to deliver 0.5 μl of virus bilaterally into the VTA (AP: ±1.0 mm, ML: ±1.0 mm, DV: - 8.5 mm) [37]. The following combination of viruses were co-infused: 166 nl of GAD1-Cre-AAV 2/10 + 333 nl of Ef1α-DIO-hM3D-AAV 2/10 (GABA targeted hM3D), or 166 nl GAD1-Cre-AAV 2/10 + 333 nl CMV-FLEX-tdTomato-AAV 2/10 (GABA targeted tdTomato, Figure 1A). The GAD1-Cre-AAV2/10 expresses Cre only in neurons that express glutamate decarboxylase 1. Co-infusion results in hM3D expression in GAD1+ GABAergic neurons [34]. All rats were given carprofen (5 mg/kg) to manage post-operative pain for two days and then an additional 3 days of recovery.

**Figure 1.**
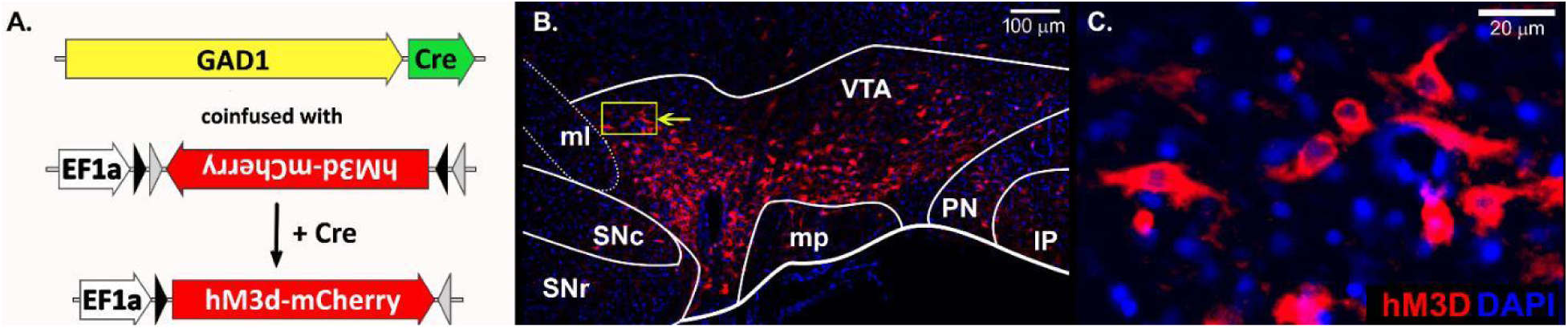
Combinatorial viral targeting of activating DREADDs to VTA GABA neurons. **(A)** AAV viral constructs that when co-infused allow for restriction of hM3D-mCherry DREADD to GAD1+ neurons. **(B)** Spread of virus within VTA (hM3D-mCherry (red) with DAPI counterstain (blue)), yellow box and arrow indicate area magnified to the right **(C)**.

### 2.3. Drugs

Clozapine N-oxide (CNO, NIDA Drug Supply Program, Bethesda, MD) was dissolved in 0.9% sterile saline solution to a final concentration of 0.3 mg/ml or 1 mg/ml CNO and administered at 1 ml/kg of body weight i.p. 30 minutes before test sessions.

### 2.4. Testing Apparatus

FI experiments were performed in eight operant self-administration chambers (30 cm x 29 cm x 23 cm, Med-Associates Inc., Georgia, VT), fitted with two nose poke ports surrounding a center reward cup containing a fluid dispenser and housed in larger sound isolation boxes. A cue light was placed directly above the dispenser and a speaker which emitted a 5 kHz tone was situated on the opposite wall. Chambers were also equipped with a house light in the upper center panel on the wall opposite the cue light and reward cup. Locomotor activity experiments used six open field chambers (43.2 cm x 43.2 cm x 30.5 cm, Med-Associates Inc., Georgia, VT) equipped with three 16-beam infrared photo arrays.

### 2.5. Procedures

#### 2.5.1. Acquisition of Sucrose Self-administration

Rats were trained to nose poke for 10% sucrose on a fixed ratio 1 schedule (FR1; Figure 2A). At the session start the house light and active nose poke light would turn on and remain on throughout the duration of the session. A response made in the active nose poke was reinforced by delivery of 70 µl 10% sucrose solution and activation of a cue light above the liquid dispenser over 4 seconds. Nose pokes made in the active port during the sucrose delivery, or at any time in the inactive nose poke, were recorded but had no programmed consequences. Sessions were 1 hr in length, or until 120 reinforcers were earned, whichever occurred first, and acquisition criteria was ≥100 reinforcers across two consecutive sessions.

**Figure 2.**
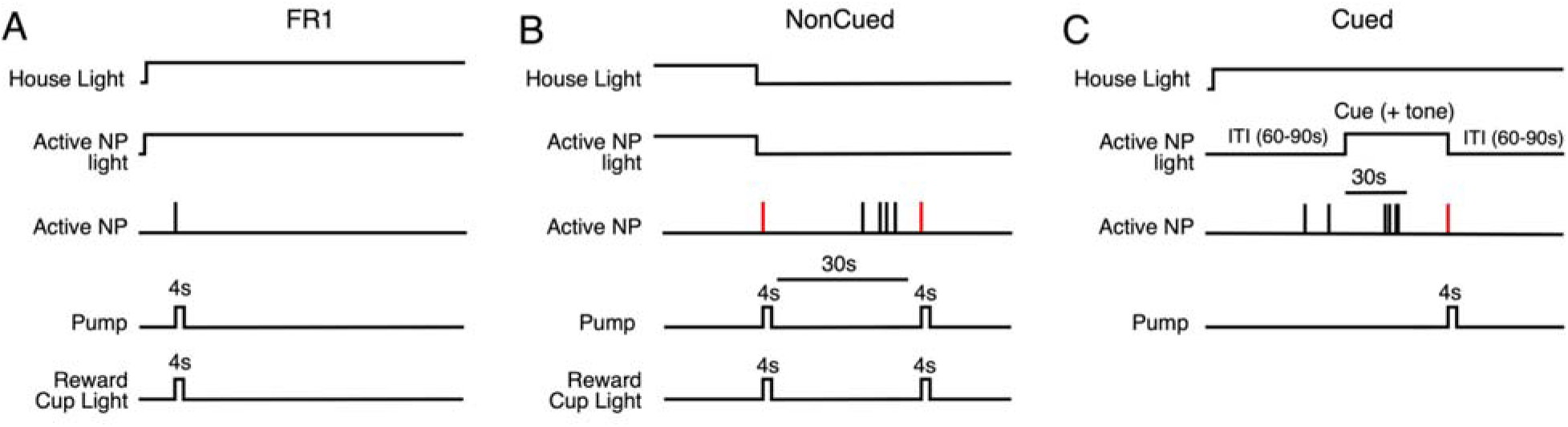
Fixed Ratio and Fixed Interval training schematics. **(A)** FR1: activation of the house light and active nose poke light signaled the start of FR session, and remained on throughout the session. Every nose poke in the active hole was reinforced except during reinforcer delivery. (**B)** Non-Cued FI: Activation of the house light and active nose poke light signaled the start of the session but both lights went out for the remainder of the session. Every response in the active nose poke port made after the 30s fixed interval was reinforced. (**C)** Cued FI: Activation of the house light signaled the start of cued FI30 session, and remained on throughout the session. The ITI preceded a tone that signaled the start of the 30s FI.

#### 2.5.2. Fixed Interval, Non-Cued

After acquisition of sucrose self-administration, the reward contingencies were changed to a non-cued FI 30s (Figure 2B), modified from Emmons, *et al*. [38]. All aspects of the FI30 Non-Cued schedule were the same as the FR schedule except that after the first response in the active nose poke port, the house light and active nose poke light turned off and remained off throughout the duration of the session. A 30s fixed-interval was imposed before the next reinforcer could be earned. Responses made during sucrose delivery, the 30s interval, or in the inactive nose poke port were recorded but had no programmed consequences. Sessions lasted two hrs or until 100 rewards were obtained, whichever occurred first.

#### 2.5.3. Fixed Interval, Cued

The FI30 Cued procedure differed from the FI30 Non-Cued, in the following ways: 1) an inter-trial interval (ITI, 60-90s) occurred between the 30s fixed intervals, 2) the inclusion of a 5 kHz tone during the 30s fixed interval until a rewarded response was made, 3) the house light remained on throughout the duration of the session, 4) the active nose poke light was not illuminated at any time, and 5) the cue light above the liquid dispenser did not illuminate during reinforcer delivery (Figure 2C). Any responses made during the ITI, 30s fixed interval, or on the inactive nose poke hole were recorded but had no programmed consequence. Sessions lasted for 2 hrs, or until 100 rewards were obtained, whichever occurred first.

### 2.6. Locomotion

All rats were tested for locomotor activity after the completion of FI testing. Rats were habituated to the locomotor chamber 1 hr per day for two days. The following day rats were placed in the chamber for 30 minutes, removed to inject either sterile saline (0.9%) or CNO (0.3 mg/kg, i.p.) and then immediately returned to the chamber for 65 minutes. Data from the first five minutes after the injection were omitted from the statistical analysis. Data are presented as the mean ± SEM in 5-minute bins.

### 2.7. Histology

After all behavioral tests, rats were anesthetized with sodium pentobarbital (390 mg/kg, i.p.) and transcardially perfused with 0.1M phosphate-buffered saline followed by 10% formalin. Brains were then placed in formalin at 4°C overnight, transferred to 30% sucrose solution until sunk, and then sectioned to 35 µm, obtained using a sliding microtome (American Optical Company, Buffalo, NY, model No. 860) and stored in 0.1M PBS + 0.02% sodium azide in 4°C until examination with a Leica TCS SP8 epifluorescent microscope.

### 2.6. Statistical Analysis

All statistics were calculated using GraphPad Prism 8 (GraphPad Software, San Diego, CA), Statistica 13 (TIBCO Software, Inc., Palo Alto, CA), and Microsoft Excel for Office 2016 (Redmond, WA); graphs were made with GraphPad Prism 8, MATLAB (MathWorks, Natick, MA), and raster plots were made in Python programming language. Response time histograms were normalized to the highest average response time to investigate changes in timing apart from changes in response rate [19]. All data were analyzed by two-way mixed design analysis of variance (ANOVA) and multiple comparison test (Tukey’s HSD) analyses were performed when significant interactions between factors were detected. In one case, a test of simple effects was used to further evaluate an interaction that trended toward significance. Magnitude of treatment effect size was reported as eta-squared, (η^2^). The level of significance was set to α < 0.05.

## III. Results

### 3.1. Verification of viral placements

Specificity of our combinatorial viral targeting system has been previously verified by immunohistochemistry and electrophysiology [34]. We verified the viral placements by histological examination of native fluorescence, which revealed robust viral expression diffused throughout the VTA (Figure 1B, 1C).

### 3.2. Non-Cued Fixed Interval 30s

Responding in FI schedules is characterized by an increase in frequency as the end of the interval becomes imminent, resulting in a “scalloping” pattern [18, 39], clearly seen in the cumulative event records after vehicle (VEH) treatment (blue line, Fig. 3A and B from one representative rat). The open circles indicate a rewarded response. Chemogenetic activation of VTA GABA with 0.3 and 1 mg/kg CNO disrupted this pattern (red and green lines, Figure 3A and B). These data are also presented as raster plots (first 50 trials, Figure 4). Each row represents an individual 30s Non-Cued FI trial, where responding during the FI is shown as black vertical tick marks (within the first 30s), and ends when the rewarded response is emitted (red vertical tick marks, >30-90s). Under VEH conditions, the majority of responding started 15-20 seconds into the 30 second interval, and most rewards were earned within 5 seconds after the end of the 30s interval, indicating precise temporal control (Figure 4A). However, CNO treatment reduced unrewarded responses during the 30s interval, and rewards were earned much later after the 30s interval ended, up to a minute or more after the end of interval (Figure 4B and C). The mean ± SEM response time histograms from VTA GABA tdTomato and hM3D rats are presented in Figure 5A and B, respectively. Under VEH conditions, or after CNO treatment in tdTomato control rats, the highest ratio of responses was emitted in the last 2 seconds of the Non-Cued 30s FI. Additionally, most rewards were earned in the five seconds after the end of the 30s FI. However, treatment with CNO resulted in less responses during the Non-Cued FI, and increased the time to rewarded responses after the 30s FI elapsed. We next determined the area under the curve (AUC) for individual histograms. To more precisely identify the effect of VTA GABA activation on responding, the AUC during the 30s FI and the 60s immediately after the end of the FI were analyzed separately. A two-way ANOVA on the AUC of the Non-Cued FI (Figure 5C) revealed a significant effect of hM3D or tdTomato *virus*; *F*_1, 14_ = 5.11, *p* = 0.04, η^2^ = 0.15), no significant effect of CNO or VEH *drug pretreatment* (*F*_2, 28_ < 1) and no significant interaction of *virus x drug pretreatment* (*F*_2, 28_ = 1.92, *p* = 0.16, η^2^ = 0.02). A slight increase in responding during the FI was seen in rats with hM3D virus expression, however pretreatment with CNO increased responding in rats with hM3D once the Non-Cued FI had ended. A two-way ANOVA on the 60s after the Non-Cued FI AUC (Figure 5D) revealed a significant effect of *virus* (*F*_1, 14_ = 6.76, *p* = 0.021, η^2^ = 0.15), a significant effect of *drug pretreatment* (*F*_2, 28_ = 6.62, *p* = 0.004, η^2^ = 0.11) and a significant interaction of *virus x drug pretreatment* (*F*_2, 28_ = 11.14, *p* < 0.001, η^2^ = 0.19). Multiple comparison tests showed an increase in AUC post-FI after pretreatment with 0.3 mg/kg (*p* < 0.001) and 1 mg/kg (*p* < 0.001) CNO in the hM3D groups when compared to both VEH controls. Together, these data indicate that the majority of the difference in AUC can be attributed to increases in the time to obtain the reinforcer at the end of the Non-Cued 30s FI.

**Figure 3.**
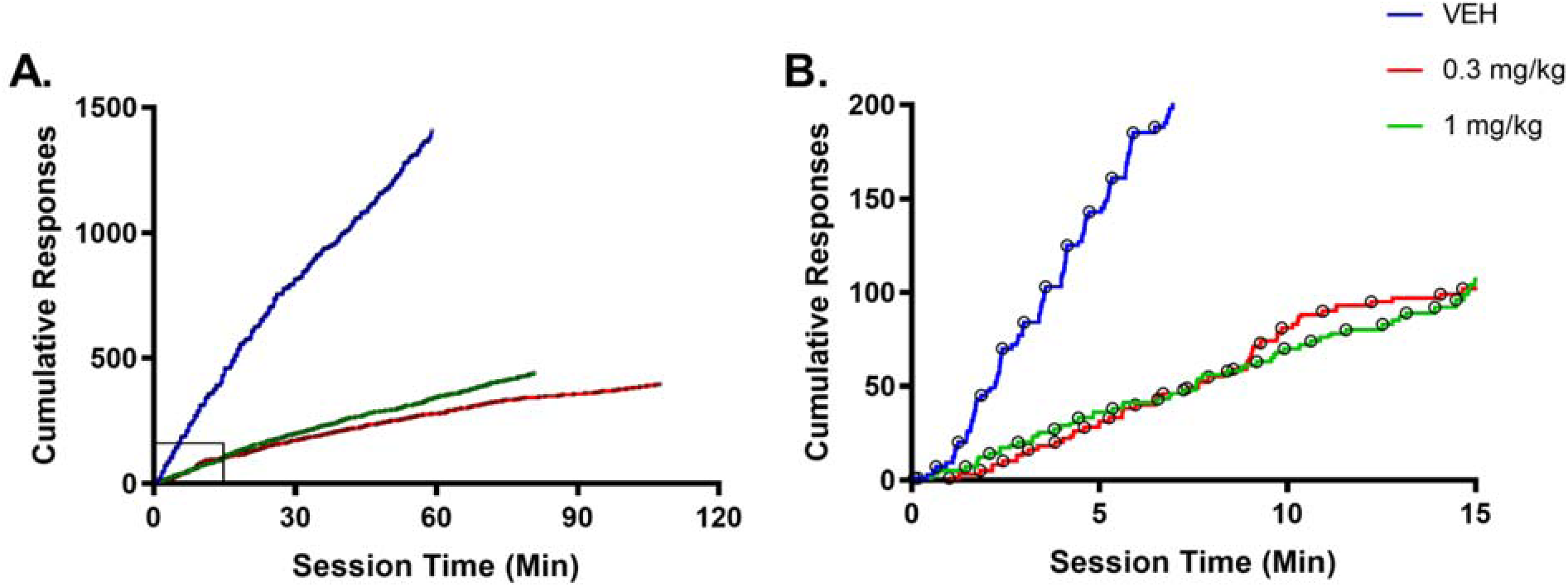
VTA GABA activation disrupts responding in a Non-Cued fixed interval task. An individual cumulative event record after i.p. vehicle (VEH) or CNO (0.3 mg/kg or 1 mg/kg). **(A)** Cumulative event records for the entire session; the black box is the area shown in **(B)**, the first 15 min of the session reveal overall rate was slower, and scalloping was disrupted. Open circles represent rewarded responses.

**Figure 4.**
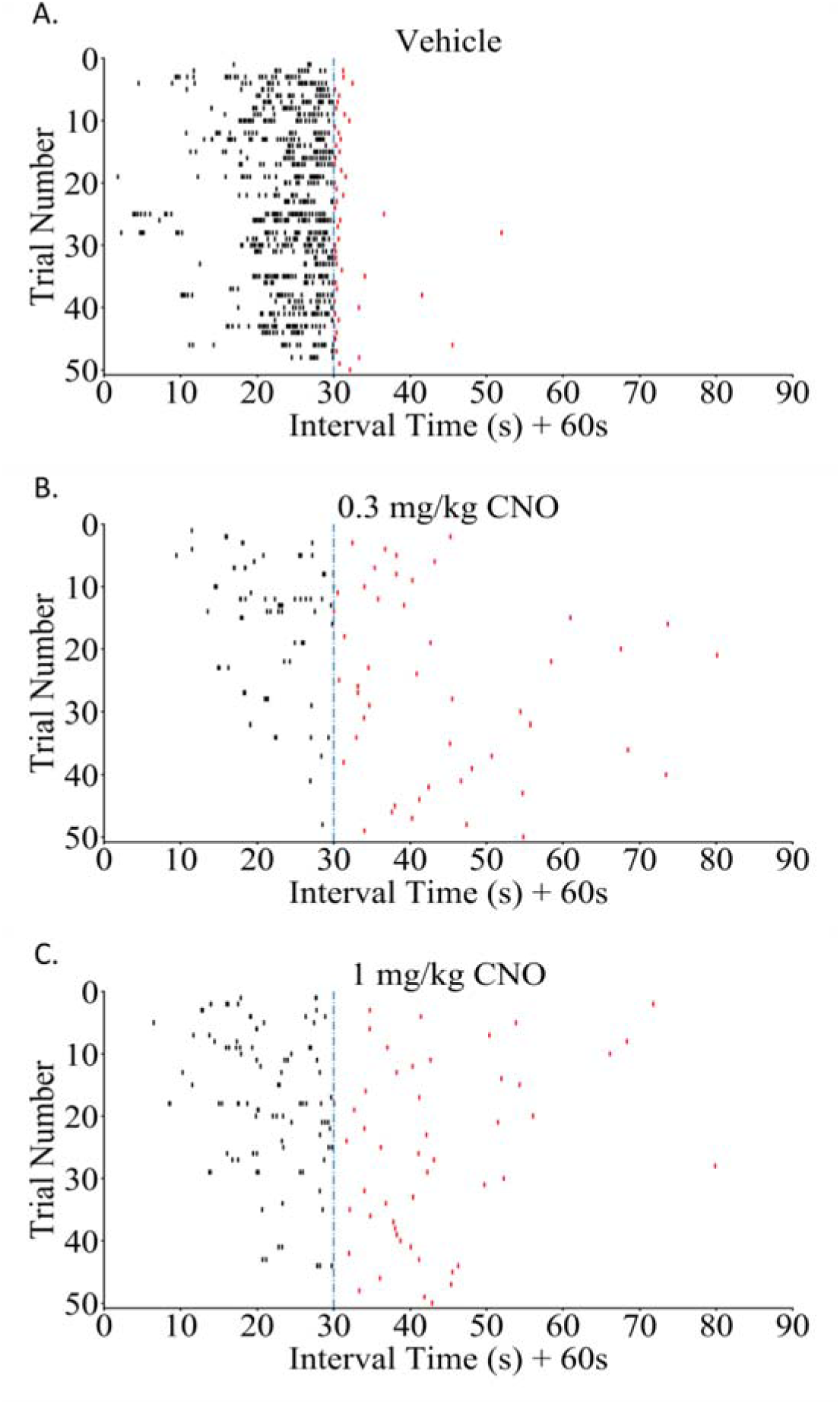
Raster plots reveal VTA GABA activation results in rats responding less during the 30s interval, and taking longer to obtain the reward when the interval is not cued. Representative raster plot from one individual for **(A)** VEH, **(B)** 0.3 mg/kg CNO, or **(C)** 1 mg/kg CNO. The data are derived from the same individual in Figure 3. Only first 50 trials are shown. Each line across the abscissa represents a single trial in the session. Responses made within the 60 additional seconds after the interval ended represent rewarded nose pokes.

**Figure 5.**
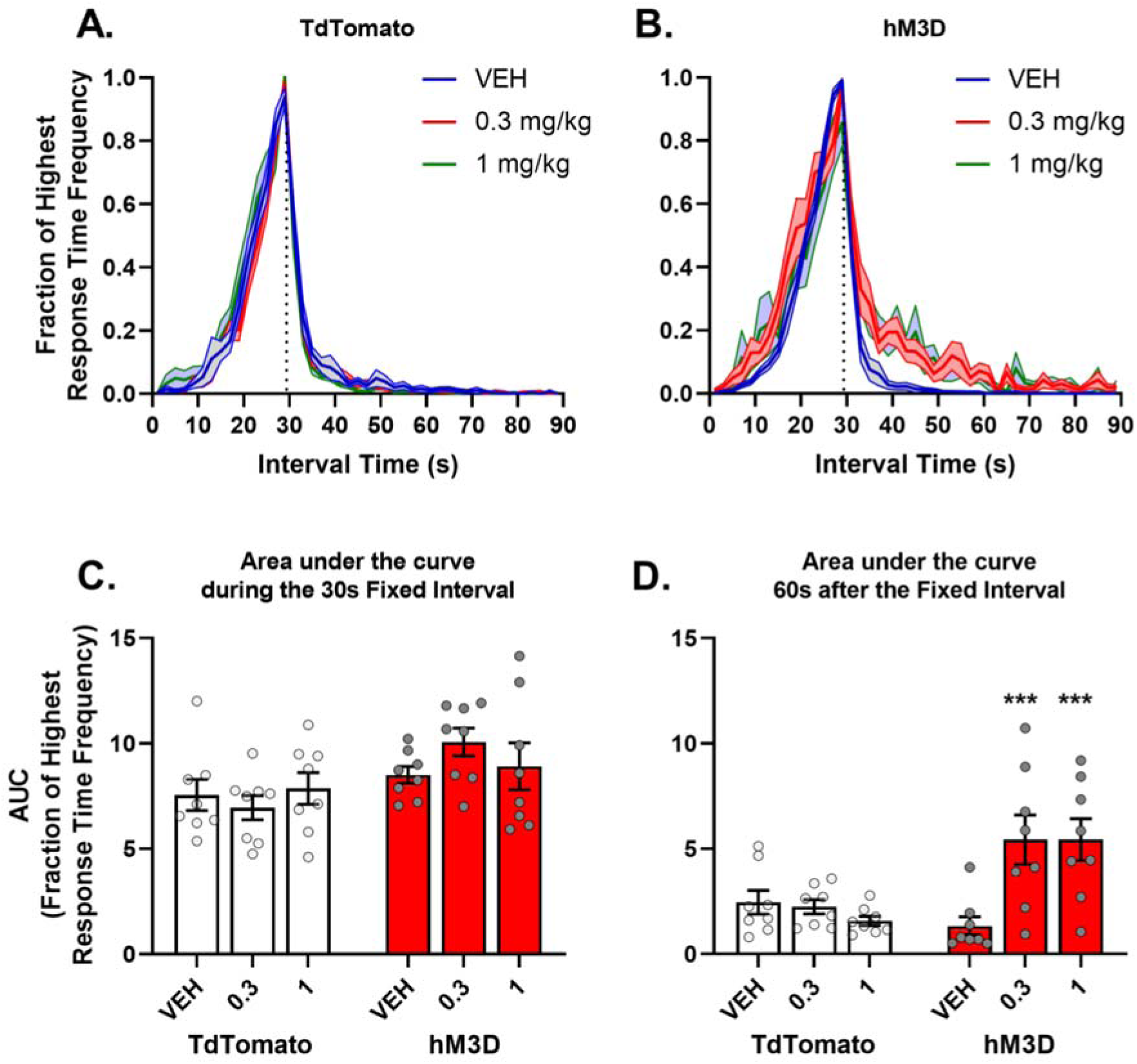
VTA GABA activation disrupts temporal control in a FI30 Non-Cued task. Response time histograms for **(A)** TdTomato and **(B)** hM3D were normalized to the highest average response. AUC for **(C)** 30s FI, and **(D)** 60s post FI. The results in are expressed as means ± SEM. Asterisks represent significant differences of either 0.3 or 1 mg/kg CNO (i.p.) compared to VEH control (*p<0.05, **p<0.01, ***p<0.005). All groups *n* = 8.

We also analyzed the latency to emit the first response after the last earned reward, which marked the end of the post-reinforcement pause and the start of responding for the end of the next interval (Figure 6A). Indeed, pretreatment with CNO increased the latency to nose poke after reinforcer delivery. A two-way ANOVA revealed no significant effect of *virus* (*F*_1, 14_ < 1), a significant effect of *drug pretreatment* (*F*_2, 28_ = 5.41, *p* = 0.01, η^2^ = 0.15) and a significant interaction of *virus x drug pretreatment* (*F*_2, 28_ = 4.60, *p =* 0.019, η^2^ = 0.12). Multiple comparison tests in hM3D rats showed an increase in the length of the average post-reinforcement pause after 0.3 mg (*p =* 0.003) and 1 mg/kg (*p =* 0.037) groups when compared to their VEH control.

**Figure 6.**
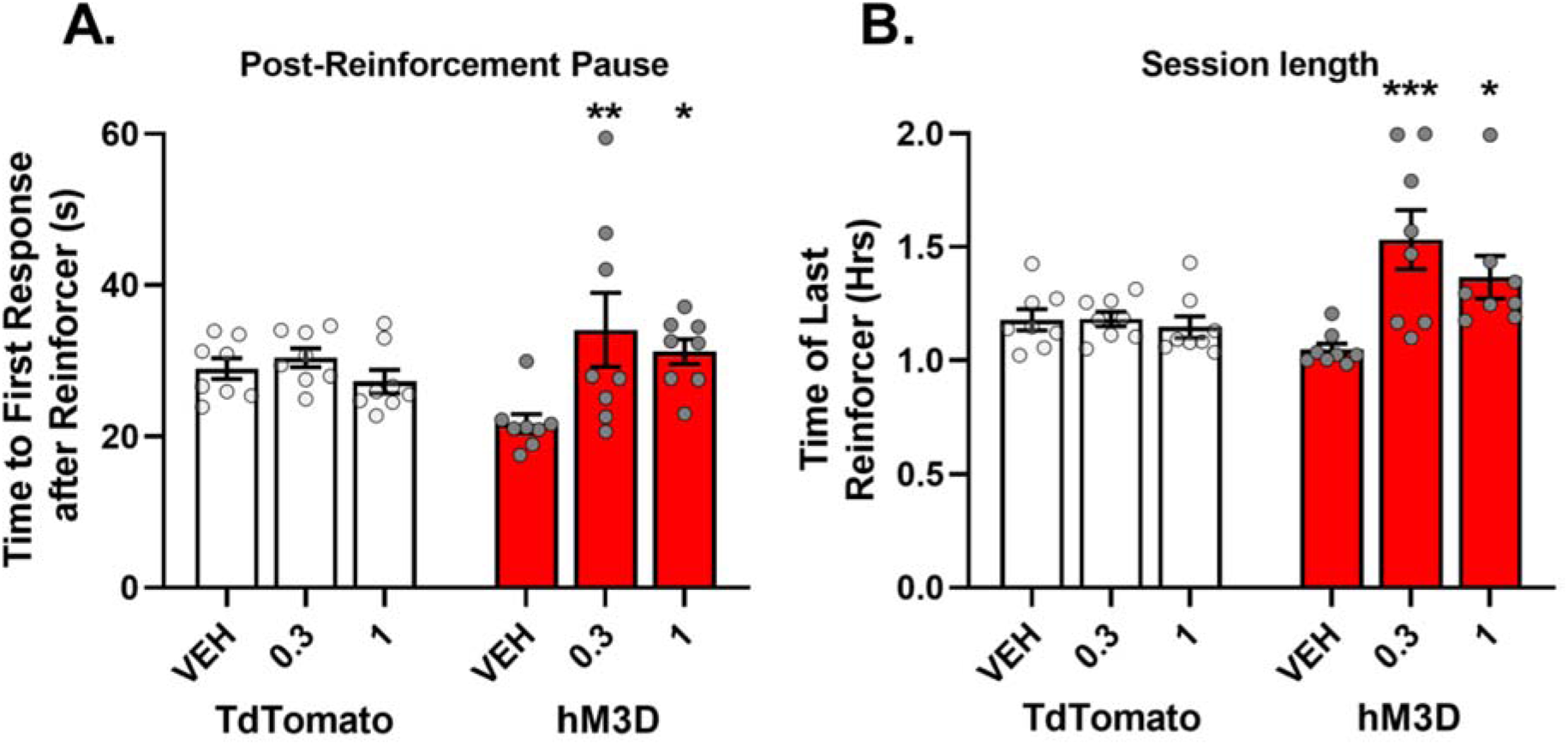
Post-reinforcement pause and session length are increased after VTA GABA activation. **(A)** Time to first response after a reinforcer, and **(B)** time of last reinforcer during Non-Cued FI. The results in are expressed as means ± SEM. Asterisks represent significant differences of either 0.3 or 1 mg/kg CNO (i.p.) compared to VEH control (*p<0.05, **p<0.01, ***p<0.005). All groups *n* = 8.

We also evaluated the session length after rats reached the 100-reinforcer cutoff. CNO pretreatment increased the session length by 1.45x and 1.30x after 0.3 and 1.0 mg/kg CNO, respectively (Figure 6B). The increased length appeared to arise from the rats taking longer to respond to obtain the reinforcer after the 30s FI had elapsed. A two-way ANOVA of the time when the last reward was delivered revealed no significant effect of *virus* (*F*_1, 14_ =4.05, *p* = 0.064, η^2^ = 0.08), a significant effect of *drug pretreatment* (*F*_2, 28_ = 7.31, *p* = 0.004, η^2^ = 0.16) and a significant interaction of *virus x drug pretreatment* (*F*_2, 28_ = 7.56, *p =* 0.002, η^2^ = 0.16). Multiple comparison tests in hM3D rats showed an increase in session time after pretreatment with 0.3 mg/kg (*p* < 0.001) or 1 mg/kg (*p =* 0.017) CNO compared to their VEH control. No effect was seen in tdTomato controls.

### 3.2. Cued FI30

Previous data from our lab showed that VTA GABA activation disrupted responding to reward-predictive cues [34]. Therefore, in this study we determined how VTA GABA activation influences behavior during intervals whose start is marked with reward-predictive cues. Rats in the Cued FI30 group were trained in a fixed interval 30s schedule of reinforcement where the start of the FI30 was signaled by the onset of a tone, which continued until the reinforcer was obtained. Thus, this cue marked the beginning of the FI but not the end. The inclusion of the tone and ITIs decreased the total trial time, and no subjects reached the 100 reinforcer cutoff before the end of the 2-hr session. VTA GABA activation disrupted the pattern of responding as indicated by a lack of scalloping in the cumulative event record from a representative rat (0.3 and 1.0 mg/kg CNO, versus VEH, Figure 7A and B). The cumulative response rate curves under VEH conditions appear to be less steep than those in the Non-Cued condition, most likely a result from the ITIs. Raster plots (first 50 trials, Figure 8) once again demonstrate the dysregulation of responding in a representative rat after receiving CNO. Pretreatment of hM3D rats with CNO disrupted the frequency of responding during and after the end of the interval (Figure 9A and B). To determine the nature of this disruption, the average AUCs during the Cued FI and after the FI were analyzed similar to the Non-Cued FI task and as seen in Figure 5. A two-way ANOVA on the Cued 30s FI AUC (Figure 9C) revealed no significant effect of *virus* (*F*_1, 14_ <1), no significant effect of *drug pretreatment* (*F*_2, 28_ = 2.67, *p* = 0.087, η^2^ = 0.09) and no significant interaction of *virus x drug pretreatment* (*F*_2, 28_ = 2.36, *p* = 0.11, η^2^ = 0.08). Pretreatment with CNO increased the AUC after the end of the interval. A two-way ANOVA of the AUC of the 60s after the fixed interval (Figure 9D) revealed no significant effect of *virus* (*F*_1, 14_ = 2.13, *p* = 0.17, η^2^ = 0.07), a significant effect of *drug pretreatment* (*F*_2, 28_ = 9.55, *p* = 0.002, η^2^ = 0.15) and a significant interaction of *virus x drug pretreatment* (*F*_2, 28_ = 9.12, *p* < 0.001, η^2^ = 0.14). Multiple comparison tests showed an increase in AUC post-FI after pretreatment with 0.3 mg/kg (*p* = 0.0002) and 1 mg/kg (*p* = 0.0004) CNO in the hM3D groups when compared to VEH. Together these data indicate that the majority of the difference in AUC can be attributed to increased time to emit the rewarded response after the Cued 30s FI. We also observed that VTA GABA activation produced a decrease in the total number of responses emitted, reinforcers earned and an increase in responses during the ITI (Figure 10). A two-way ANOVA on total responses made in a session (Figure10A) revealed no significant effect of *virus*, a significant effect of *drug pretreatment* (*F*_2, 28_ = 3.8, *p* = 0.036, η^2^ = 0.10) and no significant interaction of *virus x drug pretreatment* (*F*_2, 28_ = 1.91, *p =* 0.17, η^2^ = 0.08). The lack of an interaction on this metric was surprising; therefore, to further probe the nature of this result, simple effect tests on the impact of CNO at each level of virus treatment were conducted. Regardless of CNO dose, tdTomato control rats showed no change in total responding (*F*_2, 28_ < 1, not significant), however hM3D rats demonstrated less responding as a function of CNO dose (*F*_2, 28_ = 5.24, *p* = 0.011). Multiple comparison tests revealed the nature of this difference to be dose-dependent with less total responding between rats pretreated with 1 mg/kg when compared to VEH controls (*p* = 0.018). Regarding the effect on total number of reinforcers earned (Figure 10B), a two-way ANOVA on total reinforcers earned revealed a significant effect of *virus* (*F*_1, 14_ = 5.70, *p* = 0.032, η^2^ = 0.14), no significant effect of *drug pretreatment* (*F*_2, 28_ = 3.07, *p* = 0.062, η^2^ = 0.08) and a significant interaction of *virus x drug pretreatment* (*F*_2, 28_ = 4.21, *p =* 0.25, η^2^ = 0.11). Multiple comparison tests revealed a decrease in total number of reinforcers earned in hM3D rats with a pretreatment of 0.3 mg/kg (*p* = 0.017) and 1 mg/kg (*p* = 0.004) when compared to the VEH controls.

**Figure 7.**
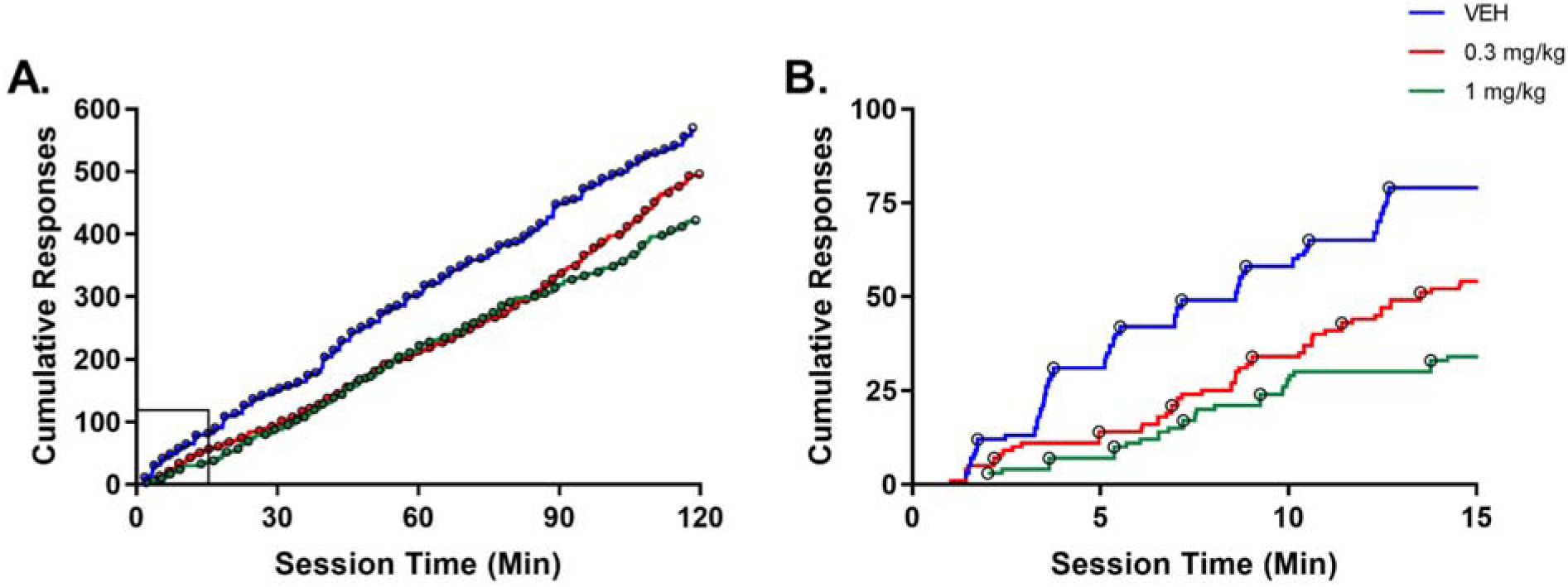
VTA GABA activation disrupts responding in a cued fixed interval task. An individual cumulative event record after i.p. VEH or CNO (0.3 mg/kg or 1 mg/kg). **(A)** Cumulative event records for the entire session; the black box is the area shown in **(B)**, the first 15 min of the session. Open circles represent rewarded responses.

**Figure 8.**
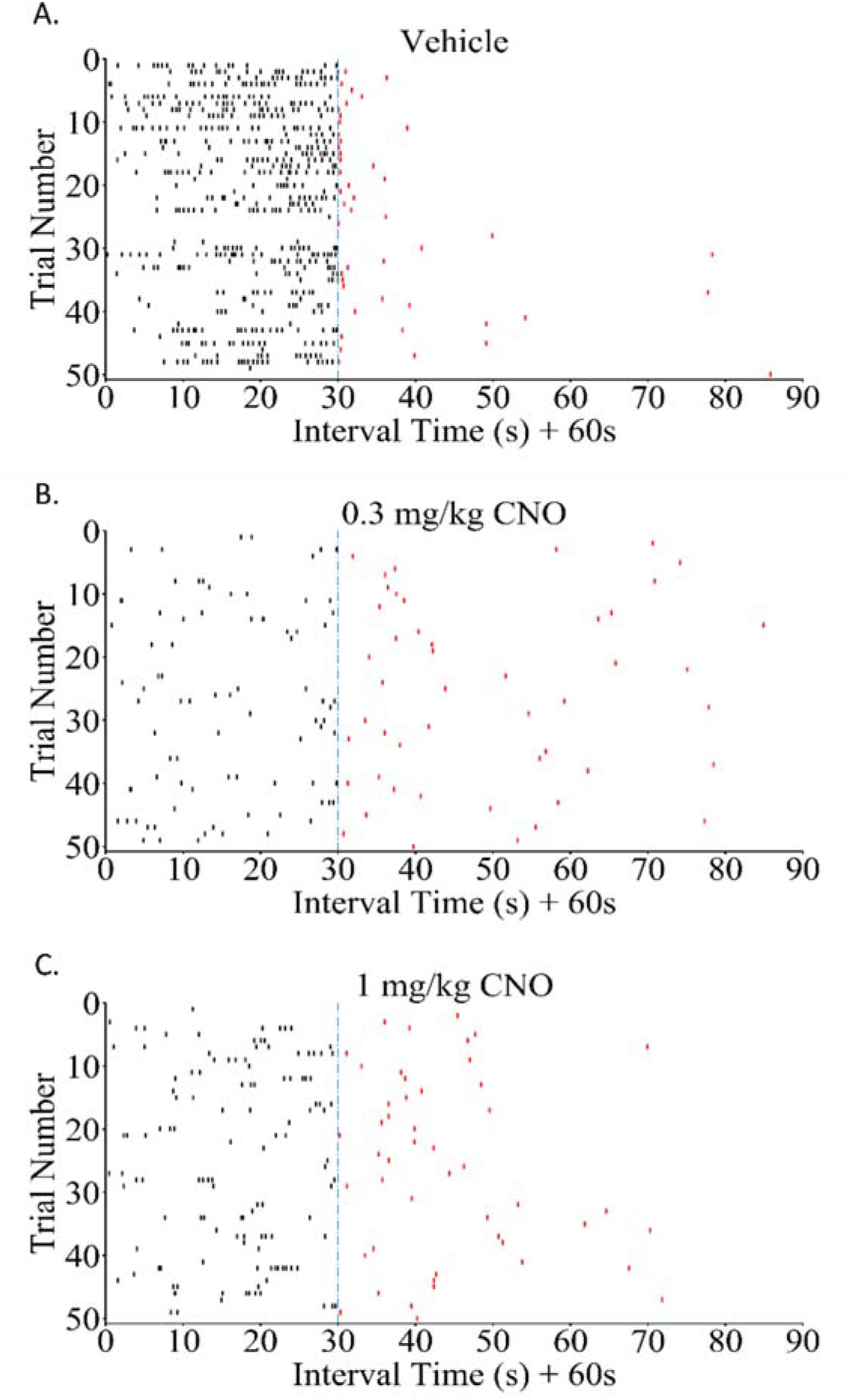
Raster plots reveal VTA GABA activation results in rats responding less during the 30s interval, and taking longer to obtain the reward when the interval is cued. Representative raster plot for **(A)** VEH, **(B)** 0.3 mg/kg CNO, and **(C)** 1 mg/kg CNO. Same rat as in Figure 7. Each line across the abscissa represents a single trial in the session. Responses made within the 60 additional seconds after the interval ended represent rewarded nose pokes.

**Figure 9.**
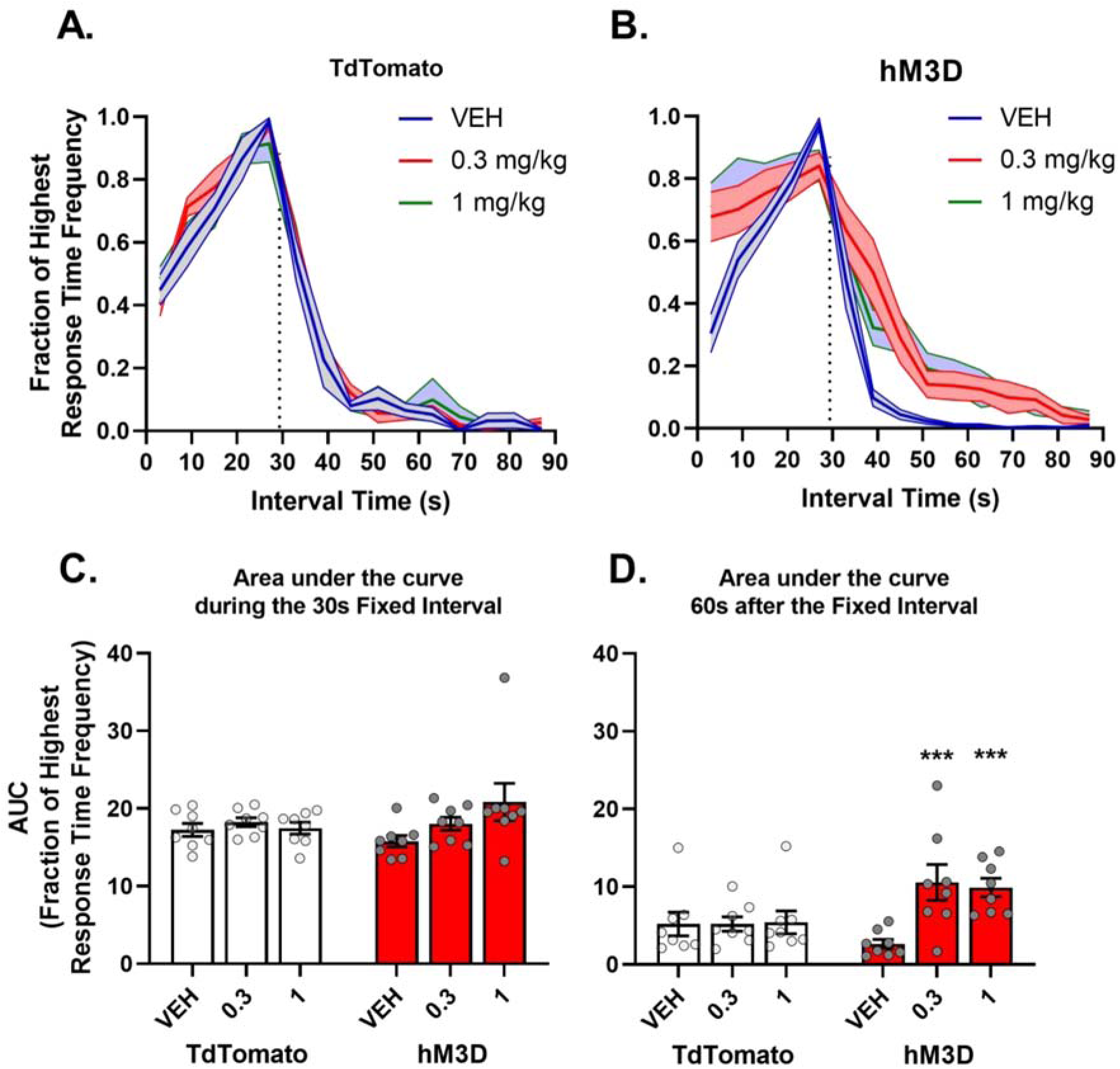
Response times reveal VTA GABA activation disrupts temporal control in a FI30 Cued task. Response time histograms for **(A)** TdTomato and **(B)** hM3D were normalized to the highest average response. AUC for **(C)** 30s FI, and **(D)** 60s post FI. The results in are expressed as means ± SEM. Asterisks represent significant differences of either 0.3 or 1 mg/kg CNO (i.p.) compared to VEH control. (*p<0.05, **p<0.01, ***p<0.005). All groups *n* = 8.

**Figure 10.**
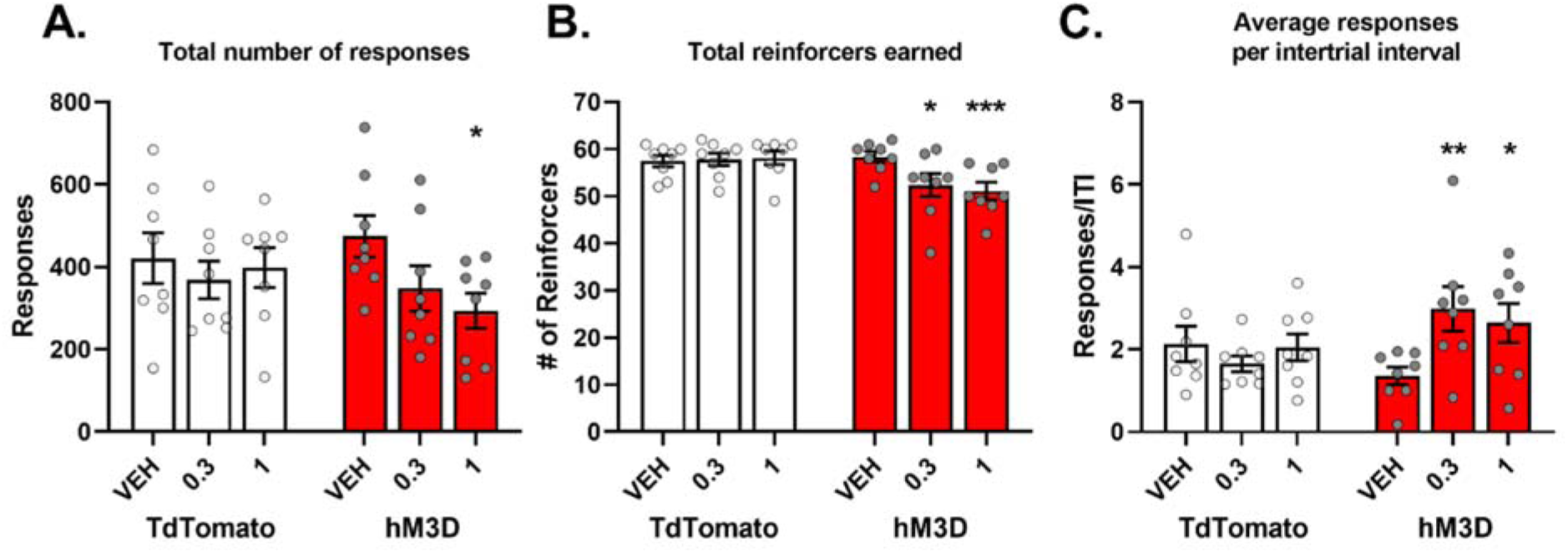
Responses, reinforcers, and responding during ITI are disrupted after VTA GABA activation. **(A)** Total responses made in the session. (**B)** Total reinforcers earned in the session. **(C)** The responses per intertrial interval. The results in are expressed as means ± SEM. Asterisks represent significant difference of either 0.3 or 1 mg/kg CNO (i.p.) compared to VEH control (*p<0.05, **p<0.01, ***p<0.005). All groups n = 8.

There was no effect in tdTomato controls. Finally, the frequency of responding during the inter-trial interval was evaluated and a two-way ANOVA (Figure 10C) revealed no significant effect of *virus* (*F*_1, 14_ < 1), no significant effect of *drug pretreatment* (*F*_2, 28_ = 2.45, *p* = 0.105, η^2^ = 0.06) and a significant interaction of *virus x drug pretreatment* (*F*_2, 28_ = 6.12, *p =* 0.006, η^2^ = 0.14). Multiple comparison tests revealed an increase in responding in hM3D rats with a pretreatment of 0.3 mg/kg (*p* = 0.002) and 1 mg/kg (*p* = 0.016) when compared to the VEH controls.

### 3.3. Locomotion

We also tested rats in an open field to determine if any of the effects seen on operant responding may arise from VTA GABA alterations of general locomotor activity (Figure 11). A two-way ANOVA on the first 30 minutes prior to VEH or /CNO pretreatment (Figure 11B) revealed no significant main effect of *pretreatment group* (*F*_1, 15_ = 1.73, *p =* 0.201), *virus* (*F*_1, 15_ = 1.25, *p =* 0.306) nor a significant *pretreatment group x virus* interaction (*F*_1, 12_ < 1, not significant), indicating that the rats had similar levels of baseline habituation locomotor activity. Two-way ANOVA of the last 60 minutes after drug treatment (Figure 11C) revealed no significant main effects of *drug pretreatment* (*F*_1, 15_ < 1), *virus* (*F*_1, 15_ = 4.09, *p =* 0.061) nor a significant *virus x drug pretreatment* interaction (*F*_1, 12_ < 1), further indicating that VTA GABA activation CNO did not alter locomotor activity.

**Figure 11.**
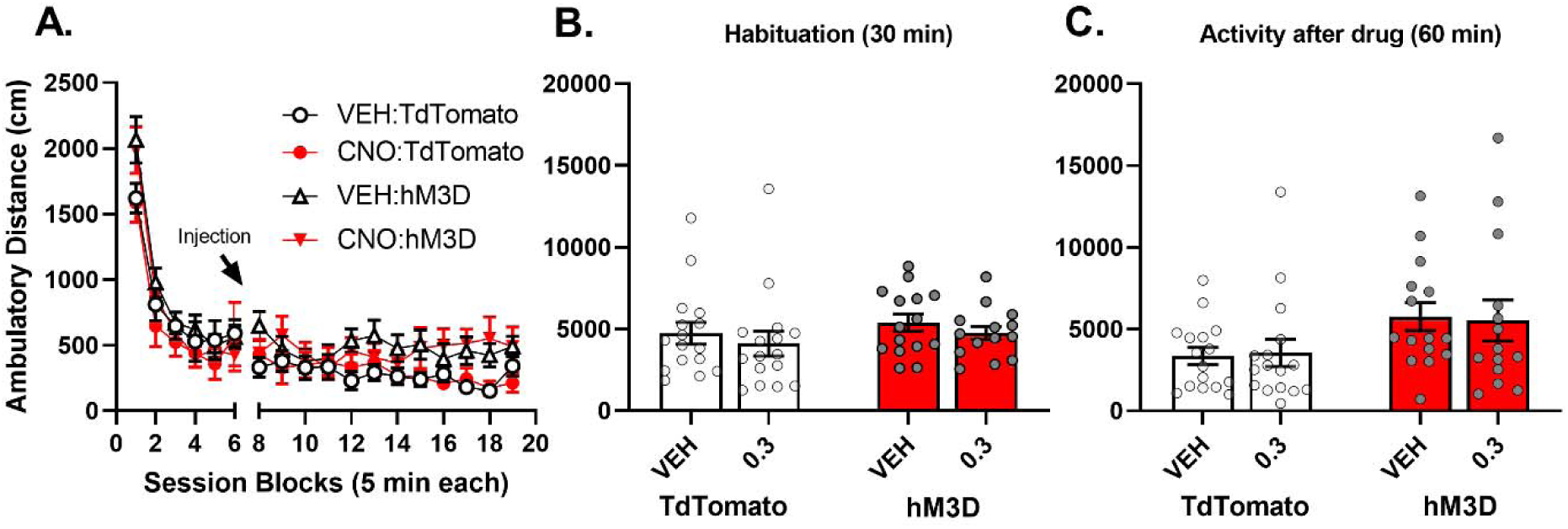
Locomotor activity was not changed after VTA GABA activation. **(A)** Locomotor activity was measured for 95 minutes total in which VEH or CNO was administered 30 minutes after being placed in an open field chamber rats. **(B)** Locomotion over the habituation period of 30 minutes and **(C)** the remaining time in the session. The results in are expressed as means ± SEM. *All groups* n *= 8*.

## IV. Discussion

Dopamine influences a subject’s internal clock [21, 23, 24], and disruption of DA transmission can alter the perception of intervals of time. For example, disease states or drugs that disrupt DA function (e.g. Parkinson’s disease, dopamine antagonists) slow the internal clock, and time is perceived as passing more quickly as evidenced by an “overestimation” of the interval time (see [40] and [21] for review). In contrast, conditions that enhance dopamine function (e.g. schizophrenia, dopamine agonists) speed up the internal clock, and time is perceived as passing more slowly. Elegant work done by Cheer and colleagues has demonstrated that NAc dopamine levels decrease as the end of a fixed interval approaches, contributing to an acceleration of responding and the traditional scalloping pattern observed with FI schedules [18]. Further, increasing dopamine in the NAc using the cannabinoid receptor 1 agonist WIN 55 212-2 resulted in premature responding during a FI30s schedule.

Given the relationship between mesocorticolimbic DA and interval timing, we hypothesized that activation of VTA GABA would disrupt dopamine neurotransmission resulting in delayed responding in a FI30s schedule of reinforcement. Indeed, our data show that in the Non-Cued FI task, nose poke responses were maximal at the end of the FI, (i.e., the 30s time point) after vehicle pretreatment (Fig. 5B), but VTA GABA activation shifted the pattern of responding, such that responding was distributed throughout the entire FI compared to controls. CNO pretreated hM3D animals would often obtain the same total number of reinforcers as controls, but did so more slowly, often overshooting the FI entirely (e.g. the first response was emitted after the 30s FI elapsed). When responding during the FI did occur, the scalloping was less pronounced and elongated, although the normalized histogram still showed some responding towards the end of the 30s FI.

Recently we have shown that chemogenetic activation of VTA GABA neurons reduces responding to cues that have acquired incentive motivational strength in an operant model [34]. The inclusion of a predictive cue creates a contingency between the reinforcer and the cue that precedes it, which may speed the acquisition and accuracy of responding [41]. Thus, we predicted that VTA GABA activation would produce a greater disruption of interval timing when the operant response was also supported by reward predictive cues. We therefore repeated these experiments in a Cued FI30 schedule, where the start of a FI is marked by an auditory cue that sounds until the reward is delivered [42, 43]. In the Cued FI task, VTA GABA activation produced an entirely different pattern of responding than controls, which was characterized by a complete flattening of the distribution of responses throughout the FI. Specifically, vehicle pretreated rats demonstrated a typical scalloped distribution of responses with greater frequency of responding towards the end of the FI (Fig. 7A and B, right panel, blue line). In contrast, VTA GABA activation resulted in a nearly absent scallop, (Fig. 7A and B red and green line), reinforced responses were emitted much later after the FI had elapsed compared to controls, and there was an increase in responding during the ITI. Importantly, this disrupted pattern of responding during the Cued FI task was profoundly different than that observed in the Non-Cued FI task, in which the basic scalloped distribution of responses was retained even after CNO pretreatment. Additionally, VTA GABA activation resulted in a decrease in the total number of reinforcers obtained in the Cued FI task (Fig. 10B).

While previous studies examining timing saw leftward shifts in responding during the FI after increasing DA transmission [18], or rightward shifts after decreasing DA transmission [38], our results revealed that VTA GABA activation completely disrupted temporal control with no discernible pattern in the Cued FI task. Since reward seeking is a multivariate process [42], with dopamine influencing many aspects, it is important to recognize that manipulating VTA GABA neurons likely produces a complex behavioral phenotype, as multiple aspects of reinforcement and cue processing are likely affected. The more pronounced behavioral disruption in the Cued FI task may speak to the mechanisms by which VTA GABA activation are affecting operant responding in general. Previous results in the lab have demonstrated that VTA GABA activation disrupts responding to sucrose predictive incentive cues [34]. It seems likely that rats trained with a cue indicating the start of an FI engage dopamine-based attention and incentive salience processes associated with the ability of reward predictive cues to regulate their responses. Since phasic DA peaks are known to shift to the earliest predictive cue from the food reinforcer they predict [44-46], if the tone in the Cued FI task induced DA signaling, then VTA GABA activation might disrupt this process. In contrast, performance on the Non-Cued FI30 schedule does not require responses to predictive cues. Thus, our results with the Non-Cued FI30 schedule would rely more on GABA’s effects on the internal clock of the animal, which is regulated by DA release upon reward delivery and consumption. Notably, the changes we observed in both FI schedules, including the slower rate of responding, do not appear to result from a disruption in overall locomotor activity, suggesting that the effects seen here were likely not related to arousal.

Chemogenetic VTA GABA activation in our study should decrease both local and long-range DA transmission. The PFC also receives long-range DA input, and is critical for top-down processing of motivated behaviors, including controlling the temporal aspects of reward-seeking. For example, D_1_ receptor antagonism in the PFC decreases responses anticipating the interval end in a fixed interval 20s timing task, while overall prefrontal cortex inactivation disrupts temporal control all together [19], similar to our results from the Cued FI task. While our current results can be mechanistically explained by local VTA GABA induced decreases in DA firing and decreased NAc DA, future studies should assess the role of VTA GABA projections, such as those to the NAc and prefrontal cortex, in influencing interval timing. In addition, lesions in the substantia nigra (SN) and the neighboring striatonigral circuit also lead to the complete loss of temporal control [19, 47]. Indeed, Parkinson’s patients have a disrupted sense of timing which is restored after L-DOPA treatment [22, 23]. Future studies determining the role of GABA regulation in this the SN should also be undertaken.

In summary, chemogenetic VTA GABA activation disrupts temporal processing of interval timing in a manner consistent with a decrease in DA output throughout the mesocorticolimbic DA circuit. VTA GABA activation decreases the ability of rats to respond to reward predictive cues in operant schedules by disrupting the ability of rats to perform the correct behavioral sequence in response to reward predictive cues [34] as well as the ability to correctly anticipate the timing of operant responding.

## V. Acknowledgements

We thank Dr. Kumar Narayanan and Ben DeCorte for their continuous help and insight into fixed interval procedures, Thomas Bassett and Li Li for their technical help. This work was supported by The Whitehall Foundation (C.E.B), and R21 DA043190 (C.E.B). The funding sources had no involvement in the conduct of the research or preparation of the article.

